# Airway basal stem cells are necessary for the maintenance of functional intraepithelial airway macrophages

**DOI:** 10.1101/2024.06.25.600501

**Authors:** Tristan Kooistra, Borja Saez, Marly Roche, Alejandro Egea-Zorrilla, Dongzhu Li, Dilanjan Anketell, Nhan Nguyen, Jorge Villoria, Jacob Gillis, Eva Petri, Laura Vera, Zuriñe Blasco-Iturri, Neal P. Smith, Jehan Alladina, Yanting Zhang, Vladimir Vinarsky, Manjunatha Shivaraju, Susan L. Sheng, Meryem Gonzalez-Celeiro, Hongmei Mou, Avinash Waghray, Brian Lin, Azadeh Paksa, Kilangsungla Yanger, Purushotama Rao Tata, Rui Zhao, Benjamin Causton, Javier J. Zulueta, Felipe Prosper, Josalyn L. Cho, Alexandra-Chloe Villani, Adam Haber, Jayaraj Rajagopal, Benjamin D. Medoff, Ana Pardo-Saganta

## Abstract

Adult stem cells play a crucial role in tissue homeostasis and repair through multiple mechanisms. In addition to being able to replace aged or damaged cells, stem cells provide signals that contribute to the maintenance and function of neighboring cells. In the lung, airway basal stem cells also produce cytokines and chemokines in response to inhaled irritants, allergens, and pathogens, which affect specific immune cell populations and shape the nature of the immune response. However, direct cell-to-cell signaling through contact between airway basal stem cells and immune cells has not been demonstrated. Recently, a unique population of intraepithelial airway macrophages (IAMs) has been identified in the murine trachea. Here, we demonstrate that IAMs require Notch signaling from airway basal stem cells for maintenance of their differentiated state and function. Furthermore, we demonstrate that Notch signaling between airway basal stem cells and IAMs is required for antigen-induced allergic inflammation only in the trachea where the basal stem cells are located whereas allergic responses in distal lung tissues are preserved consistent with a local circuit linking stem cells to proximate immune cells. Finally, we demonstrate that IAM-like cells are present in human conducting airways and that these cells display Notch activation, mirroring their murine counterparts. Since diverse lung stem cells have recently been identified and localized to specific anatomic niches along the proximodistal axis of the respiratory tree, we hypothesize that the direct functional coupling of local stem cell-mediated regeneration and immune responses permits a compartmentalized inflammatory response.

## INTRODUCTION

Airway epithelial basal stem cells are multipotent stem cells that are essential for the long-term maintenance of the other airway epithelial cell types^1–3^. It has been demonstrated that basal cells signal to their own daughter cells by direct cell-to-cell contact through Notch signaling, effectively serving as niches for their progeny^1,2^. Basal cells are also known to mediate inflammation via the secretion of chemokines and cytokines ^4–12^, and reciprocally, the immune system is known to modulate stem cell activity in multiple organs, including the lung^8,9,13^.

In recent years, the murine tracheal epithelium has received considerable attention because the overall structure of the murine trachea mirrors aspects of the epithelial cell composition of human conducting airways^14–16^. In particular, the murine trachea and proximal bronchi, but not more distal mouse airways, contain the basal stem cells that are abundant throughout the human conducting airways where the pathology of airway disorders, including asthma, are localized^17^. Thus, while most murine modeling of airway inflammation has been focused on the analysis of distal lung tissue, the murine trachea is well suited for the study of disease-relevant stem cell-to-immune cell communication.

Communication between airway epithelial cells and airway myeloid cells, such as dendritic cells (DCs) and macrophages, is critical for pulmonary immunity^4–10^. Elucidating the biological basis of this communication has led to specific therapies that target epithelial cell-derived cytokines, such as thymic stromal lymphopoietin (TSLP) and IL-33, which have been shown to have clinical efficacy in treating asthma^18,19^. However, we are only just beginning to understand the heterogeneity of airway immune cells and the mechanisms that govern airway epithelial cell and immune cell interactions^20–25^.

Airway myeloid cells within the airway are organized into a network comprised of an intraepithelial and a subepithelial compartment. Intraepithelial myeloid cells physically contact airway epithelial cells, and it is thought that they participate in immunity by directly surveilling the airway lumen^26,27^. Recently, a unique population of intraepithelial airway macrophages (IAMs) has been identified in the trachea and proximal bronchi of mice^13,25^. These cells are the predominant MHC II^+^ myeloid cell population within the murine tracheal epithelium and seem to be important for airway epithelial repair following injury^13^. In addition, there is evidence that these cells can take up inhaled antigens and present these antigens to airway-resident T cells ^25^.

Herein, we show that depletion of airway basal stem cells functionally disrupts the local network of IAMs and thereby alters antigen-driven inflammatory responses. Interestingly, the responses in the distal airways of the mouse, which lack basal stem cells, remain unaffected, suggesting that the stem cell effect is local. Indeed, we show that the compartmentalized nature of this phenomenon is explained by the necessity for contact-mediated signaling in which basal cell-derived Dll1 activates Notch receptors on IAMs. Furthermore, IAMs require tonic Notch signaling from basal cells to maintain MHC II expression and for proliferation. Notch blockade in IAMs not only decreases MHC II expression but is also associated with reduced allergic airway inflammation in a murine model of asthma. Interestingly, human counterparts to IAMs also exhibit ongoing Notch activation and thus may participate in pathologic epithelial-immune crosstalk in airway disease.

## RESULTS

### A unique intraepithelial airway macrophage population

A large body of work has previously described a network of dendritiform airway myeloid cells within the airways that were thought to be DCs^4–7,27^. However, a recent study exploring the immune composition of the murine trachea identified a population of intraepithelial airway macrophages (IAMs) within the upper airway epithelial layer with a similar dendritiform morphology^13^. Using a combination of single-cell RNA sequencing (scRNAseq) analysis^2^ and immunofluorescent staining of the trachea, we confirmed that these intraepithelial dendriform cells represent IAMs (Figure 1 and Figure S1). IAMs sit adjacent to CK5^+^ basal stem cells and demonstrate overlap with markers of conventional DC type 2 cells (cDC2), including CD11c (*Itgax*), high levels of MHC II, and signal regulatory protein α (*SIRPα*) (Figure 1A-1E, Figure S1A-S1C). However, unlike cDC2, IAMs lack expression of *Zbtb46* and express *Irf8* (Figure 1B and Figure S1A). Additionally, IAMs express macrophage-associated markers, including *Mafb*, *CX3CR1*, *F4/80 (Adgre1)*, *CD68*, and complement-associated genes, including *C1q* and *C5ar1* (Figure 1B and Figure S1A-S1E). Importantly, the expression of *Scimp* and *Tgfbr1,* which has been previously associated with IAMs^13^, is also observed in this population (Figure S1A). Furthermore, lineage tracing analysis using *Zbtb46-Cre; mTmG* reporter mice that specifically label conventional dendritic cells (cDCs)^28^ does not result in the labeling of IAMs. Interestingly, we only found rare cDCs within the epithelium at homeostasis, and the vast majority of intraepithelial MHC II^+^ cells were IAMs (Figure S1F). To further characterize this unique immune population, we used a *Cx3cr1-GFP; Ccr2-RFP* dual-reporter mouse line. We found that IAMs express CX3CR1 and include a subfraction that co-expresses CCR2 (Figure 1F), suggesting that IAMs undergo homeostatic turnover and are replaced by circulating CCR2^+^ monocytic precursors, in agreement with prior data in the setting of injury^9,13^.

**Figure 1.**
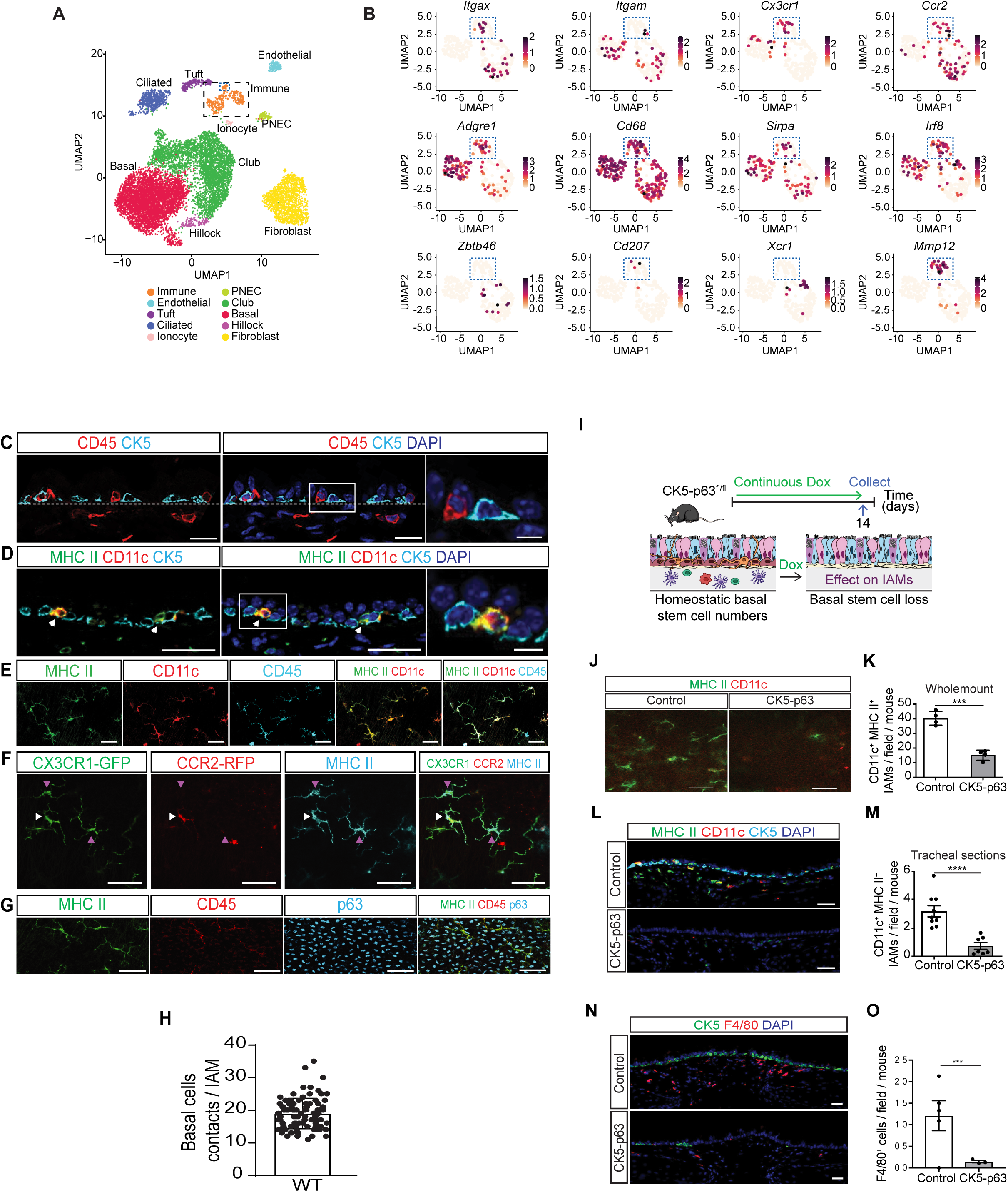
Basal stem cells are necessary for the maintenance of intraepithelial airway macrophages (IAM). **A,** UMAP clustering of mouse tracheal cells isolated from wild-type (WT) mice. Black dashed box highlights airway immune cells, with the intraepithelial airway macrophage (IAM) subset indicated by the smaller blue dashed box. **B,** Differential expression of markers of dendritic cells or macrophages in tracheal immune cells including *Itgax (CD11c), Itgam (CD11b), Cx3cr1, Ccr2, Adgre1 (F4/80), Cd68, SIRPα, Irf8, Zbtb46, Cd207 (Langerin), Xcr1* and *Mmp12*. Blue dashed box highlights IAM population. **C,** Immunostaining for basal stem cells (CK5, cyan) and immune cells (CD45, red) without (left) or with (right) DAPI (blue) on WT tracheal sections. **D,** Immunostaining for IAM markers MHC II (green) and CD11c (red), and basal stem cell marker CK5 (cyan) without (left) or with (right) DAPI (blue) on WT tracheal sections. White arrowheads point to CD11c+/MHC II+ IAMs within the epithelium. **E,** Wholemount confocal imaging for MHC II (green), CD11c (red) and CD45 (cyan) on WT tracheal explants at the epithelial plane. **F,** Wholemount immunostaining for GFP (CX3CR1, green), RFP (CCR2, red), and MHC II (cyan) on tracheal explants from *CX3CR1-GFP; CCR2-RFP* mice. White arrows point to CCR2+ IAMs within the epithelium. Purple arrows point to CX3CR1+/MHC II+ cells within the epithelium. **G,** Wholemount immunostaining for MHC II (green), CD45 (red) and basal stem cell marker p63 (cyan) on WT tracheal explants. **H,** Quantification of the number of basal stem cells (p63+) in contact with one IAM. **I,** Schematic representation of the depletion of CK5-expressing basal stem cells induced by the loss of the basal transcription factor p63 in *CK5rtTA; TetO Cre; p63^fl/fl^* mice (CK5-p63). Ciliated cells, Club cells, goblet cells, and basal stem cells are shown in blue, pink, purple, and brown respectively. Dox: doxycycline. **J,** Wholemount immunostaining for MHC II (green) and CD11c (red) in control (left) or CK5-p63 mice (right). **K,** Quantification of absolute numbers of IAMs (CD11c+/MHC II+) per 20x field in control or CK5-p63 tracheal wholemounts (n=4/group). **L,** Immunostaining for MHC II (green), CD11c (red) and CK5 (cyan) in control (upper panel) or CK5-p63 mice (lower panel). **M,** Quantification of absolute numbers of IAMs (CD11c+/MHC II+) per 20x field in control or CK5-p63 mice (n=9 control; n=7 experimental). **N,** Immunostaining for CK5 (green), and F4/80 (red) in control (upper panel) or CK5-p63 (lower panel) mouse tracheas. **O,** Quantification of absolute numbers of intraepithelial F4/80+ cells per 20x field in control or CK5-p63 mouse tracheas (n=9 control; n=7 experimental). Control and CK5-p63 mice were all treated with Dox. Experiments were repeated three times (3 independent experiments). *** *p*<0.001, **** *p*<0.0001. Data shown in the graphs are means ± SEM. Nuclei stained with DAPI (blue). Scale bar, 20μm. See also Figure S1 and S2.

These analyses also demonstrated that IAMs are localized immediately adjacent to or directly above basal stem cells (Figure 1C, 1D). Using fixed wholemount preparations of the murine trachea, we determined that each IAM is in direct, simultaneous cellular contact with 19 ± 5 p63^+^ basal stem cells on average. (Figure 1G, 1H). Thus, our findings confirm the presence of a distinct population of myeloid cells characterized by (1) an intraepithelial airway location adjacent to basal stem cells, (2) a dendritiform morphology, (3) a unique signature that aligns with a macrophage identity, and (4) high expression of MHC II, suggesting that they are antigen-presenting cells.

### Airway basal stem cells are necessary for the maintenance of functional IAMs

Due to the proximity of IAMs to basal stem cells, we sought to understand whether airway stem cells regulate adjacent IAMs. We employed *CK5rtTA; TetO-Cre; p63^fl/fl^* mice (CK5-p63 mice hereafter) to eliminate CK5-expressing airway basal stem cells from the airway epithelium by inducible deletion of p63 as previously reported^29^. p63 is the essential basal stem cell master transcription factor, which is deleted upon doxycycline (Dox) administration in drinking water in these mice, leading to basal stem cell loss without significant toxicity (Figure 1I)^29^. Consistent with this, tracheas from Dox-treated CK5-p63 mice display a decrease in epithelial thickness due to the paucity of basal cells (Figure S2A). However, a continuous layer of differentiated epithelial cells remains and displays normal expression of tight junction proteins zonula occludens-1 (ZO1), claudin-1, and occludin (Figure S2B, S2C). We confirmed basal stem cell loss in Dox-treated CK5-p63 mice compared to Dox-treated *p63^fl/fl^* control mice with p63 and CK5 staining (67±18 *vs*. 501±94 p63^+^ cells per tracheal section, and 160±20 *vs*. 663±81 CK5^+^ cells per tracheal section respectively), while club cell and FoxJ1^+^ ciliated cell numbers were unchanged (Figure S2D-S2F). Interestingly, we observed a slight increase in acetylated tubulin^+^ ciliated cells, suggesting precocious differentiation following p63 deletion in basal stem cells as previously reported^29^. Multiple rare cell types have been characterized in the tracheal epithelium with potential immunomodulatory roles^2,30–33^, and we verified that the numbers of tuft cells and ionocytes were unchanged by analyzing tracheal sections (Figure S2G-S2J) and wholemount tracheal preparations (Figure S2K-S2P) following depletion of basal stem cells. Neuroendocrine (NE) cell numbers were increased in CK5-p63 tracheas compared to control tracheas when quantified on tracheal sections (Figure S2J). However, such a difference was not observed on wholemount tracheal preparations (Figure S2K).

The analysis of intraepithelial CD11c^+^/MHC II^+^ cells (IAMs) on tracheal wholemount preparations revealed that IAMs were decreased in basal stem cell-depleted tracheas compared to controls (15.1±1.3 *vs.* 40±6.6 cells per field respectively) (Figure 1J, 1K) with similar results observed in sectioned tracheas (0.72±0.2 *vs.* 3.2±0.6 cells per field respectively) (Figure 1L, 1M). Staining for other markers expressed by IAMs confirmed our observations. Notably, quantification of F4/80^+^ cells and SIRPα^+^/MHC II^+^ cells revealed a reduction in CK5-p63 tracheas (Figure 1N, 1O, S2Q). To confirm that the loss of IAMs was not due to systemic effects of doxycycline delivered by drinking water, we administered nebulized Dox to CK5-p63 mice (Figure S2R). This method restricts tet-inducible Cre activation to the mouse airways and lung^34^. Consistent with the prior data, we again observed fewer CD11c^+^/MHC II^+^ IAMs in basal stem cell-depleted tracheas compared to controls (Figure S2S, S2T).

### Tonic Notch activation is required for the maintenance of a functional network of IAMs

To probe the mechanisms by which basal stem cells interact with IAMs, we considered that Notch signaling may play an important role given the high degree of cell-cell contact between IAMs and airway basal stem cells (Figure 1C-1E, 1G, 1H). Prior studies have implicated the Notch pathway in the development of several myeloid cell subsets^35–38^, and we have previously shown that basal stem cell-derived Notch signals directly regulate neighboring epithelial cells^1^. Based on this evidence, we hypothesized that Notch ligands from basal stem cells may signal to IAMs. To test our hypothesis, we crossed a *CD11c-mCherry* reporter mouse line with the Notch signaling *CBF:H2B-Venus* nuclear reporter mouse line to assess canonical Notch signaling activity. We found that IAMs displayed Notch signaling activity identified by nuclear reporter signal (Figure 2A). More specifically, we detected Notch2 intracellular domain (N2ICD) (Figure 2B and Figure S3A) and Notch3 intracellular domain (N3ICD) (Figure 2C and Figure S3B) by immunofluorescent staining in the nuclei of IAMs, suggesting tonic Notch2 and Notch3 activation at homeostasis.

**Figure 2.**
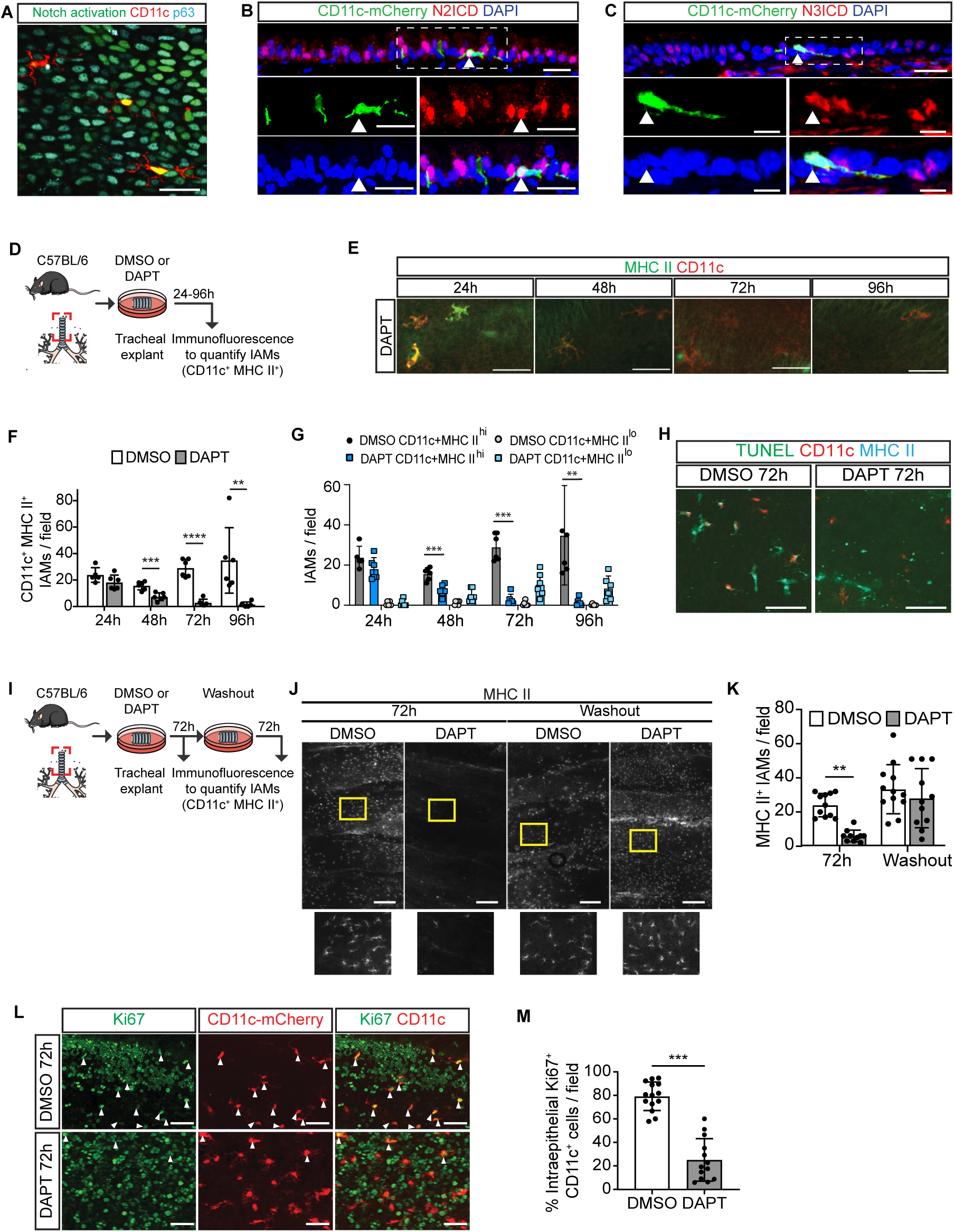
Notch activity is required for IAM functional markers and proliferation. **A,** Wholemount staining of a tracheal explant from *CD11c-mCherry; NotchCBF-H2B Venus* mice for p63 (cyan) imaged in the epithelial plane. mCherry (CD11c, red), Venus (Notch activity, green). **B,** Immunostaining for Notch2 intracellular domain (N2ICD, red) and mCherry (CD11c, green) on *CD11c-mCherry* mouse tracheas. Arrowhead points to N2ICD+ IAMs. White box indicates the area shown at higher magnification in single-channel insets below. **C,** Immunostaining for Notch3 intracellular domain (N3ICD, red) and mCherry (CD11c, green) on *CD11c-mCherry* mouse tracheas. Arrowhead points to N3ICD+ IAMs. White box indicates the area shown at higher magnification in single-channel insets below. **D,** Schematic representation of Notch inhibition experiments on tracheal explants from WT mice after treatment with the gamma-secretase inhibitors DAPT (2.5μM) for 24h, 48h, 72h and 96h. **E,** Immunostaining for MHC II (green) and CD11c (red) on wholemounts of tracheal explants treated with DAPT at different timepoints. **F,** Quantification of the absolute numbers of IAMs per 20x field after 24h, 48h, 72h and 96h of treatment with vehicle (DMSO) or DAPT (n=5-6 images/group). **G,** Quantification of the absolute numbers of CD11c^+^ MHC II^hi^ and CD11c^+^ MHC II^lo^ cells per 20x field after 24h, 48h, 72h and 96h of treatment with vehicle (DMSO) or DAPT (n=5-6 images/group). **H,** Assessment of apoptosis by TUNEL assay (green) in IAMs (CD11c (red), MHC II (cyan)), on WT tracheal explants treated with vehicle (DMSO) or DAPT for 72h. **I,** Schematic representation of Notch inhibition on tracheal explants after treatment with DAPT (2.5μM) for 72h, followed by washout. **J,** Immunostaining for MHC II on wholemounts of tracheal explants treated with vehicle (DMSO) or DAPT for 72h (left), and 72h after washout (right). Yellow boxes indicate the area shown at higher magnification in insets below. **K,** Quantification of absolute numbers of IAMs (CD11c+/MHC II+) per 20x field in these conditions (n=10-12 images/group). **L,** Wholemount staining for proliferating cells (Ki67+, green) in *CD11c-mCherry* (red) tracheal explants treated with DMSO (upper panels) or DAPT (bottom panels) for 72h. White arrows point to proliferating (Ki67+) CD11c+ cells. **M,** Quantification of the percent of proliferating IAMs (Ki67+ CD11c+) per field treated with DMSO or DAPT for 72h (n=13-14 images/group). Experiments *in vivo* were repeated three times (3 independent experiments) and experiments *ex vivo* were repeated twice. ** *p*<0.01, *** *p*<0.001 and **** *p*<0.0001. Data shown in the graphs are means ± SEM. Nuclei stained with DAPI (blue). Scale bar, 5μm (C insets), 20μm (A, B, C, E, K), 50μm (L), 200μm (I). See also Figure S3.

To further assess the importance of Notch signaling for IAMs, we excised tracheas from wild-type (WT) mice and treated them *ex vivo* with two different Notch inhibitors, DAPT and DBZ^39^ for up to 96h. Tracheal explants were then fixed and stained for CD11c and MHC II and analyzed using wholemount confocal microscopy (Figure 2D). Notch inhibition with DAPT or DBZ caused a dramatic decrease in IAMs identified by CD11c^+^/MHC II^+^ staining (Figure 2E, 2F and Figure S3C-S3E). This decrease was rapid and progressive, starting at 48h after treatment and reaching maximal inhibition 72h after treatment (Figure 2F, 2G). Of note, we did not detect apoptotic IAMs using TUNEL staining (Figure 2H). Interestingly, a CD11c^lo^MHC II^neg^ cell population with similar dendritiform morphology transiently appeared between 48-72 hours but was quickly lost (Figure 2G), suggesting the continual presence of IAMs with decreased expression of key markers. Interestingly, washout of DAPT treatment led to a recovery of MHC II^+^ IAMs within 72 hours (Figure 2I-2K). There are no circulating monocytes to replenish IAMs in our explant model suggesting that these cells either regain marker expression with DAPT washout and/or there is replacement of cells via proliferation. Consistent with this, Notch inhibition resulted in a significant decrease in the fraction of Ki67^+^ proliferating CD11c^+^ IAMs in the explants (Figure 2L, 2M).

To further investigate the role of Notch in IAM homeostasis *in vivo*, we used an inducible mouse line to target Notch signaling in established IAMs. Based on the reporter mouse data showing that MHC II^+^ IAMs express CX3CR1 (Figure 1F), we assessed the efficiency of the *Cx3cr1-CreER*; *Rosa26-tdTomato* reporter mouse line in labeling IAMs (Figure 3A). Following tamoxifen (Tam) administration, we confirmed that intraepithelial tdTomato^+^ cells co-express IAM markers, including MHC II and CD11c, and displayed a dendritiform morphology (Figure 3B, 3C). This reporter system resulted in the initial labeling of 80% of IAMs, which gradually declined over 24 weeks (Figure 3D). This suggests that under homeostatic conditions, IAMs are constantly replaced by circulating precursors, consistent with our previous observations of a small fraction of IAMs expressing CCR2 (Figure 1F). Having demonstrated the fidelity and efficiency of the *Cx3cr1-CreER* driver line to target IAMs, we next generated *Cx3cr1-CreER; mTmG; RBPJk^fl/fl^* (hereafter CX3CR1-RBPJk) mice in which canonical Notch signaling can be conditionally disrupted in CX3CR1^+^ IAMs. Furthermore, targeted IAMs are also lineage labeled by mGFP, allowing one to follow the fate of IAMs after they have lost canonical Notch activity following Tam administration (Figure 3E). Indeed, we confirmed that lineage labeled IAMs lacked RBPJk on tracheal sections (Figure S3F). When compared with *Cx3cr1-CreER; mTmG; RBPJk^+/+^* controls, we noted that the number of lineage-labeled IAMs per field was unchanged (21.2±2 *vs.* 24.2±1.4 Lineage^+^ IAMs per field in control tracheas) (Figure 3F, 3G), suggesting that Notch activation was dispensable for maintaining IAM tissue localization. However, there was a substantial reduction in lineage labeled IAMs that expressed MHC II (6.8±1.3 *vs.* 22.8±1.3 Lineage^+^ IAMs per field in control tracheas) (Figure 3F, 3H), suggesting that while IAMs remain present in the tissue in the absence of Notch activation, MHC II expression is reduced. We did not observe any apoptotic IAMs based on a lack of Caspase3 detection in CX3CR1-RBPJk tracheas (Figure 3I). Although the numbers of lineage-labeled IAM cells were unchanged in CX3CR1-RBPJk airways, we observed a reduction in the percentage of proliferating Ki67^+^ IAMs compared to controls (6.3±3.2% *vs.* 21.8±2.1% per field, respectively, Figure 3J, 3K), consistent with the effects of pharmacologic Notch inhibition *ex vivo* (Figure 2L, 2M). Altogether, these results suggest that Notch signaling is required for the maintenance of IAM functional differentiation (antigen presentation) and is also required for their proliferation.

**Figure 3.**
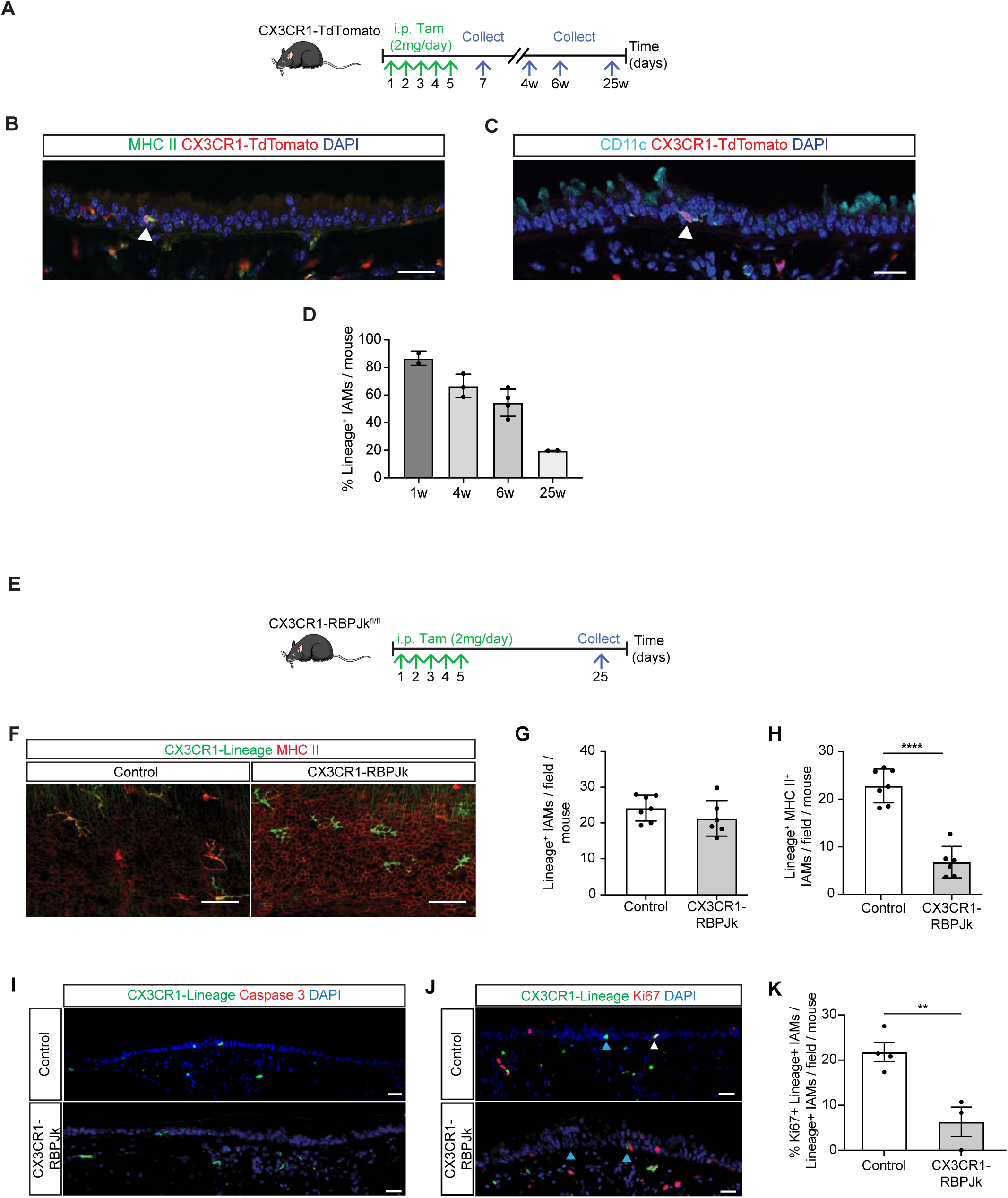
*In vivo* Notch inhibition in IAMs causes a loss of functional maturation status. **A,** Schematic representation of lineage labeling of IAMs using Tam (2mg/day) in *CX3CR1-CreER; TdTomato* mice (CX3CR1-TdTomato). i.p.: Intraperitoneal. Tam: tamoxifen. **B, C,** Immunostaining for CX3CR1-lineage labeled IAMs (TdTomato+, red) and MHC II **(B**; green) or CD11c **(C**; cyan). Arrowheads point to IAMs. **D,** Percent of CX3CR1-lineage labeled IAMs per trachea 1, 4, 6 and 25 weeks (w) after tamoxifen induction (n=2-4/group). **E,** Schematic representation of RBPJk (canonical Notch) inhibition in CX3CR1+ cells using *Cx3cr1-CreER; mTmG; RBPJk^fl/fl^* (CX3CR1-RBPJk) mice. **F,** Immunostaining for MHC II (red) and GFP (CX3CR1+, green) on tracheal wholemounts 20 days after the last tamoxifen administration. **G, H,** Quantification of absolute numbers of intraepithelial CX3CR1-lineage labeled cells (GFP+) (**G**) and double positive CX3CR1+ (GFP+) MHC II+ cells **(H)** per 20x field in control and CX3CR1-RBPJk tracheas (n=7 control, n=6 experimental). **I,** Immunostaining for CX3CR1-lineage marker GFP (green) and apoptotic marker Caspase 3 (red) in control (upper panel) or CX3CR1-RBPJk mice (lower panel) at baseline. **J,** Immunostaining for CX3CR1-lineage marker GFP (green) and proliferation marker Ki67 (red) on control (CX3CR1-TdTomato) (upper panel) or CX3CR1-RBPJk mice (lower panel) at baseline. White arrowheads point to proliferating (Ki67+) IAMs, blue arrowheads point to non-proliferating (Ki67-) IAMs. **K,** Quantification of the percentage of Ki67+ CX3CR1-lineage+ IAMs per Lineage+ IAMs per 20x field of control or CX3CR1-RBPJk tracheas (n=4 control; n=3 experimental). Experiments were repeated twice (2 independent experiments). ** *p*<0.01 and **** *p*<0.0001. Data shown in the graphs are means ± SEM. Nuclei stained with DAPI (blue). Scale bar, 20μm (B, C, I, J), 50μm (F). See also Figure S3.

### Airway basal stem cell-derived Dll1 is necessary to maintain functional IAMs

Basal stem cells are known to express high levels of the Notch ligands Jag2 and Dll1^1,39^ and have extensive contact with neighboring IAMs (Figure 1C, 1D, 1G, 1H). Thus, we hypothesized that airway basal stem cells supply a Notch ligand that activates Notch2/3 in neighboring IAMs necessary for their MHC II expression and proliferation. Using a *Dll1-mCherry* reporter mouse, we found that Dll1 is expressed in virtually all basal stem cells (Figure 4A) and in rare airway epithelial cells such as NE cells and tuft cells (Figure S3G, S3H) but not in ionocytes (Figure S3I). We then assessed the effects of deleting Dll1 from basal stem cells by employing *CK5-CreER; Rosa26-LSL-eYFP; Dll1^fl/fl^* mice (CK5-Dll1 hereafter) to remove Dll1 from CK5^+^ basal stem cells and concomitantly mark these cells with a YFP reporter following Tam administration (Figure 4B). We confirmed that YFP^+^ basal cells from Tam-treated CK5-Dll1 tracheas lacked detectable expression of Dll1 (Figure S3J) and verified that common epithelial cell populations were unchanged in frequency (Figure S3K). Tracheas from CK5-Dll1 mice displayed fewer CD11c^+^/MHC II^+^ IAMs compared to Tam-treated control mice (1.4±0.4 *vs.* 4.6±0.9 IAMs per field respectively) (Figure 4C, 4D). Furthermore, quantification on wholemount staining confirmed the reduction in CD11c^+^/MHC II^+^ IAMs following Dll1 elimination in basal stem cells compared to controls (25.1±1.8 *vs.* 39.5±2.6 IAMs per field, respectively) (Figure 4E, 4F). We also observed residual intraepithelial CD45^+^ leukocytes with similar morphology to IAMs, which lacked MHC II expression in CK5-Dll1 tracheas (Figure 4G). Finally, we targeted Jag2 expression in airway basal stem cells using *CK5-CreER; Rosa26-LSL-eYFP; Jag2^fl/fl^* mice (CK5-Jag2 hereafter) (Figure 4H). There was no difference in CD11c^+^ MHC II^+^ cells in the trachea of Tam-treated mice compared to control mice (Figure 4I, 4J), suggesting that Dll1 is the dominant basal cell-derived Notch ligand engaging IAMs.

**Figure 4.**
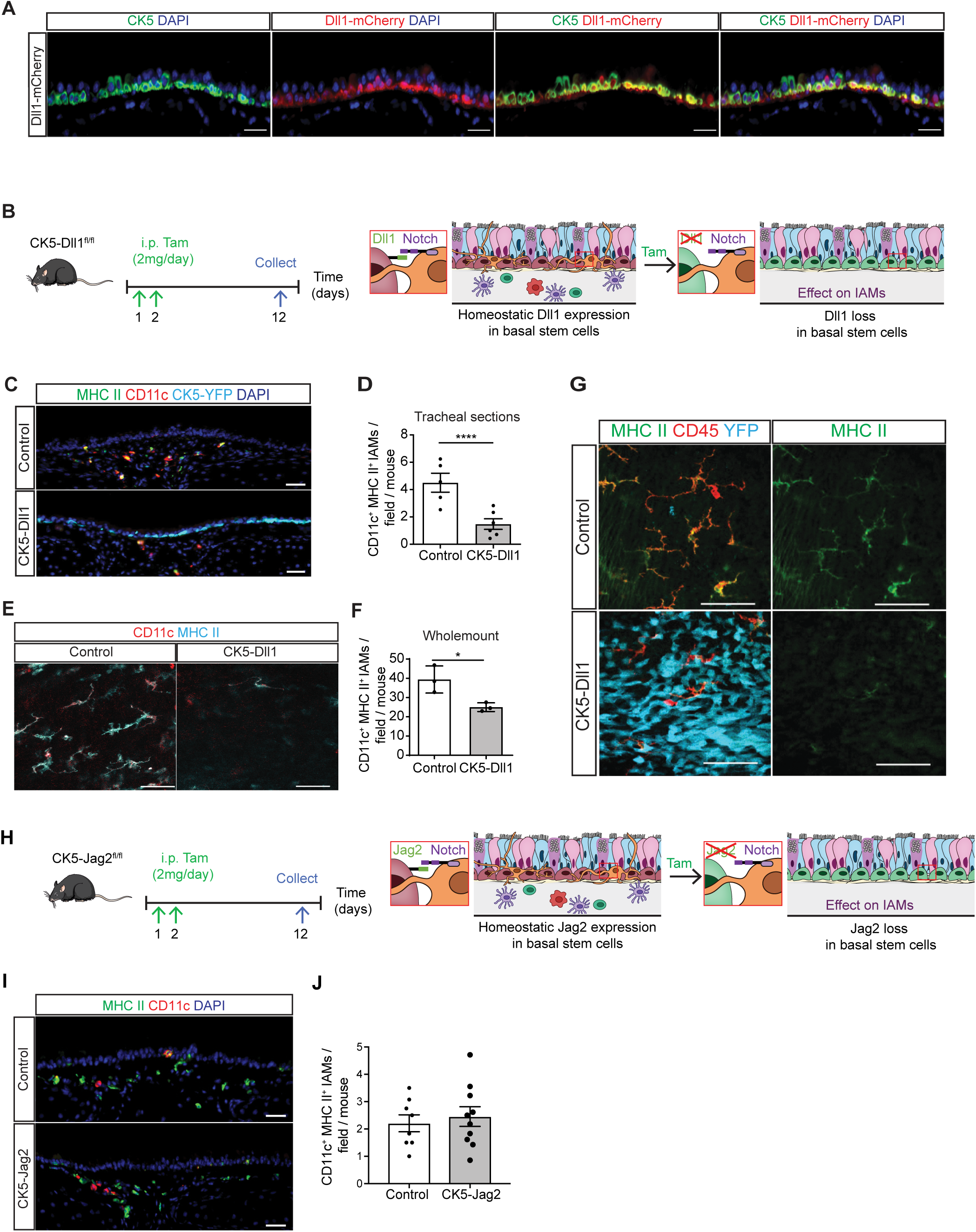
Basal stem cell-derived Notch ligand Dll1 but not Jag2 is necessary for the maintenance of IAM function. **A,** Immunostaining for Dll1 (mCherry, red) and basal stem cell CK5 (green) in *Dll1-mCherry* reporter mouse tracheas. **B,** Schematic representation of the inhibition of the Notch ligand Dll1 specifically in CK5-expressing basal stem cells using *CK5-CreER; Rosa26-LSL-eYFP; Dll1^fl/fl^*mice (CK5-Dll1^fl/fl^). i.p.: Intraperitoneal. Tam: tamoxifen. **C,** Immunostaining for CK5 lineage label YFP (cyan) and IAMs (MHC II (green) and CD11c (red)) in control (upper panel) or CK5-Dll1 mice (lower panel). **D,** Quantification of absolute numbers of IAMs (CD11c+/MHC II+) per 20x field in control or CK5-Dll1 mice (n=5 control, n=6 experimental). **E,** Wholemount staining for CD11c (red) and MHC II (cyan) on control and CK5-Dll1 tracheal explants. **F,** Quantification on wholemount system of absolute numbers of IAMs (CD11c+/MHC II+) per 20x field in control or CK5-Dll1 mice (n=3 control, n=3 experimental). **G,** Wholemount staining for CK5 lineage label YFP (cyan), CD45 (red) and MHC II (green) on control and CK5-Dll1 tracheal explants. **H,** Schematic representation of the use of *CK5-CreER; Rosa26-LSL-eYFP; Jag2^fl/fl^* (CK5-Jag2) mice to inhibit the Notch ligand Jag2 in CK5-expressing basal stem cells. **I,** Immunostaining for MHC II (green) and CD11c (red) in control (upper panels) or CK5-Jag2 mice (bottom panels). **J,** Quantification of absolute numbers of IAMs (CD11c+/MHC II+ cells) per 20x field in control or CK5-Jag2 mouse tracheas (n=8-10/group). All mice were treated with Tam. Experiments were repeated three times (3 independent experiments). * *p*<0.05, and **** *p*<0.0001. Data shown in the graphs are means ± SEM. Nuclei stained with DAPI (blue). Scale bar, 20μm (A, C, I), 50μm (E,G). See also Figure S3.

### Airway basal stem cell loss prevents allergen-induced airway inflammation

Since airway myeloid immune cells can act as sentinels, continuously sampling and processing antigens^4–7^, we assessed the functional consequences of basal stem cell deficiency on IAM using a model of airway inflammation dependent on antigen presentation. Ovalbumin (OVA) is a classic model antigen to which mice can be sensitized through intraperitoneal injection of alum-conjugated OVA. Subsequently, antigen challenge by nebulization of OVA alone results in airway antigen presentation, leading to allergic inflammation^40^ (Figure 5A). Compared to other models of allergic airway inflammation, such as the house dust mite model, the OVA model has the advantage of reduced stimulation of airway epithelial cells, innate immune responses, and pathogen recognition receptors ^41^. Thus, it is a cleaner model to look at antigen-induced inflammation in the airways. After Dox-induced deletion of p63, we again confirmed that basal cells were effectively depleted and that other epithelial cell populations remained unchanged in this context (Figure S4A, S4B) while CD11c^+^/MHC II^+^ IAMs were reduced (Figure 5B, 5C). The differentiation of mucous cells, as detected by FOXA3 and Muc5AC immunostaining as well as Alcian blue staining, was observed in OVA-sensitized and challenged control mice as expected (Figure 5D, 5E, top panels). However, there were substantially fewer newly formed mucous cells in basal stem cell-depleted tracheas after OVA sensitization and challenge (Figure 5D, 5E, bottom panels). Indeed, while control tracheas show an average of 222±44.5 Muc5AC^+^ cells per tracheal section after OVA challenge, only 73.3±16.3 mucous cells were found per tracheal section in Tam-treated CK5-p63 mice after OVA challenge (Figure 5F).

**Figure 5.**
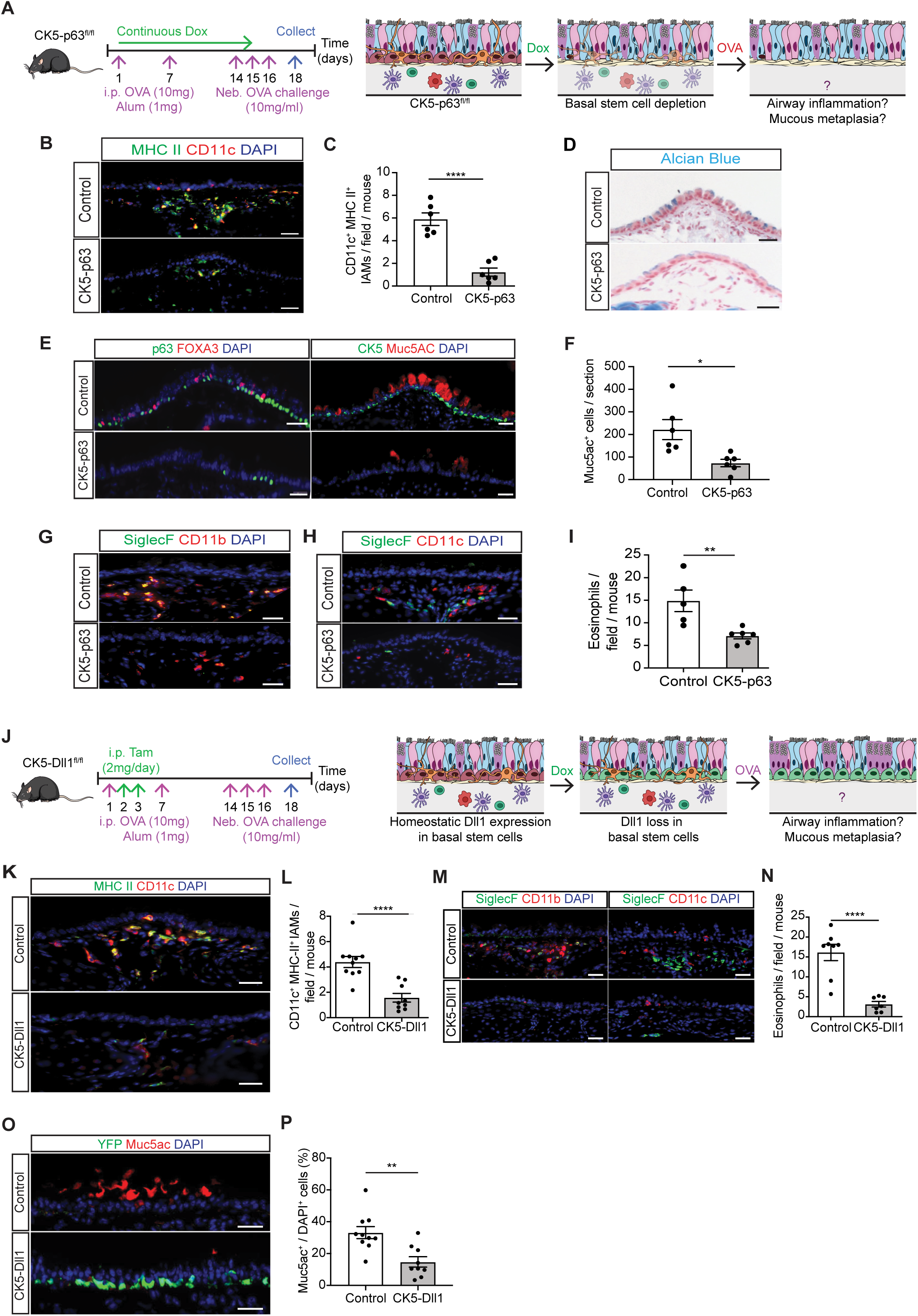
Basal stem cells are necessary for allergen-induced mucous metaplasia. **A,** Schematic representation of the depletion of CK5-expressing basal stem cells by inhibiting the basal transcription factor p63 in *CK5rtTA; TetO Cre; p63^fl/fl^* (CK5-p63) mice, and induction of an adaptive immune response leading in the development of mucous metaplasia after ovalbumin (OVA) challenges. Neb: nebulized. **B,** Immunostaining for IAMs (MHC II (green) and CD11c (red)) in OVA-treated control (upper panel) or CK5-p63 mice (lower panel) (n=4/group). **C,** Quantification of absolute numbers of IAMs (CD11c+/MHC II+) per 20x field in OVA-treated control or CK5-p63 mice (n=6/group). **D,** Alcian Blue staining of tracheas from OVA-treated control and CK5-p63 mice. **E,** Immunostaining for basal stem cells (p63, CK5; green) and mucous cells (FOXA3, Muc5AC; red) in OVA-treated control (upper panel) or CK5-p63 mice (lower panel). **F,** Quantification of Muc5AC+ cells per tracheal section in OVA-treated control and experimental mice (n=6/group). **G,** Immunostaining for eosinophils (SiglecF (green) and CD11b (red)) in OVA-treated control (upper panel) or CK5-p63 mouse tracheas (lower panel). **H,** Immunostaining for SiglecF (green) and CD11c (red) on either OVA-treated control (upper panel) or CK5-p63 mouse tracheas (lower panel) (n=6). **I,** Quantification of absolute numbers of eosinophils (SiglecF+ CD11b+ CD11c-) per 40x field in OVA-treated control or CK5-p63 mice (n=5-6/group). **J,** Schematic representation of the inhibition of the Notch ligand Dll1 specifically in CK5-expressing basal stem cells with OVA sensitization and challenge. **K,** Immunostaining for IAMs (MHC II (green) and CD11c (red)) in OVA-treated control (upper panel) or CK5-Dll1 mice (lower panel). **L,** Quantification of absolute numbers of IAMs (CD11c+/MHC II+) per 20x field of the trachea of OVA-treated control or CK5-Dll1 mice (n=10 control, n=9 experimental). **M,** Immunostaining for eosinophils (SiglecF, green) and CD11b (red) (left panels) or CD11c (red) (right panels) in OVA-treated control (upper panel) or CK5-Dll1 mice (lower panel). **N,** Quantification of absolute numbers of eosinophils (SiglecF+ CD11b+ CD11c-) per 40x field in OVA-treated control or CK5-Dll1 mice (n=10 control, n=9 experimental). **O,** Immunostaining for CK5 lineage YFP (green) and Muc5AC (red) in OVA-treated control (upper panel) or CK5-Dll1 mice (lower panel). **P,** Quantification of the percentage of Muc5AC+ mucous cells per DAPI+ cells in OVA-treated control or CK5-Dll1 mice (n=9-10/group). Control and CK5-Dll1 mice were all treated with Tam. Experiments were repeated three times (3 independent experiments). * *p*<0.05, ** *p*<0.01, and **** *p*<0.0001. Data shown in the graphs are means ± SEM. Nuclei stained with DAPI (blue). Scale bar, 20μm. See also Figure S4, S5 and S6.

We hypothesized that the lack of mucous metaplasia seen following basal stem cell loss reflected a diminished local type-2 inflammatory response^17^. Since eosinophilic infiltration is a hallmark of a type-2 immune response, we examined tracheal sections for eosinophils using a staining pattern of SiglecF^+^/CD11b^+^/CD11c^-^. We found that airway basal stem cell-depleted tracheas contained fewer eosinophils compared to controls after OVA sensitization and challenge (7.1±1.1 *vs.* 14.9±1.4 eosinophils per field, respectively; Figure 5G-5I). Of note, inflammation in the small airways and alveoli of OVA-sensitized and challenged CK5-p63 mice was comparable to control mice. Specifically, the number of eosinophils and activated CD4^+^ T cells in the lung compartment and bronchoalveolar lavage (BAL) were similar (Figure S4C-S4F), and there was no change in distal airway goblet cell formation (Figure S4G, S4H). This suggests that the loss of airway basal stem cells exclusively affects the local immune network that is adjacent to the basal stem cell niche in the murine trachea.

### Deletion of Dll1 on airway basal stem cells prevents allergen-induced airway inflammation

We next evaluated whether basal stem cell Dll1 loss would result in an aberrant response to OVA challenge (Figure 5J). Tracheas from Tam-treated CK5-Dll1 OVA-sensitized and challenged mice compared to OVA-sensitized and challenged control tracheas, displayed reduced CD11c^+^/MHC II^+^ IAM numbers (1.6±0.5 *vs.* 4.4±0.8 cells per field respectively) (Figure 5K, 5L), eosinophils (3.1±0.7 *vs.* 16.11±2 cells per field respectively) (Figure 5M, 5N), and mucous metaplasia (13.2 ± 1.6% *vs.* 34.2 ± 2.3% mucous cells per total DAPI^+^ cells in tracheas respectively) (Figure 5O, 5P). In contrast, mucous metaplasia was evident in the distal airways (Figure S4I), and no differences were observed in inflammatory cells in the BAL (Figure S4J) or tissue (Figure S4K, S4L) from OVA-sensitized and challenged CK5-Dll1 mice compared to control mice, again suggesting that the effect of altering basal stem cell signaling specifically affects the local airway immune network. In contrast, deletion of Jag2 on airway basal stem cells did not prevent mucous metaplasia following OVA sensitization and challenge (Figure S4M, S4N), consistent with the lack of effect on IAMs (Figure 4I, 4J).

We hypothesized that the observed reduction in allergic inflammation of the trachea with either airway basal stem cell depletion or basal stem cell-specific Dll1 deletion is due to the loss of IAM functional maturation. However, given the possibility that multiple immune cell types could be influenced by these basal stem perturbations, we aimed to test whether direct depletion of IAMs would phenocopy these results. We employed *Cx3cr1-CreER; R26-DTA^fl/+^* mice (CX3CR1-DTA hereafter) to deplete IAMs in the setting of OVA sensitization and challenge (Figure S5A). Intranasal 4-hydroxytamoxifen (4-OHT) administration leads to ablation of approximately half of IAMs in the trachea as measured by CD11c^+^/MHC II^+^ staining (Figure S5B). When combined with OVA sensitization and challenge, IAM depletion led to a decrease in tracheal eosinophils (386±27 *vs.* 561±76 eosinophils per mm^2^ respectively) (Figure S5C, S5D) and mucous cells (17.27 ± 2.6% *vs.* 33.8 ± 2.8% Muc5ac^+^ cells per total DAPI^+^ cells respectively) (Figure S5E, S5F) in Tam-treated CX3CR1-DTA mice compared to Tam-treated *Cx3cr1-CreER; R26-DTA^+/+^* control mice, while distal lung inflammation was preserved (Figure S5G, S5H).

Basal stem cells regulate cell fate of neighboring airway epithelial cells through the expression of Notch ligands^1^. Thus, they may directly regulate OVA-induced mucous cell differentiation independently of regulating IAM maintenance. To further assess whether the loss of basal stem cell Notch signaling might directly contribute to the loss of mucous cell metaplasia by altering the responsiveness of secretory cells to inflammation, we evaluated whether differences in goblet cell formation in CK5-p63 and CK5-Dll1 mice are related to an intrinsic altered capacity of epithelial secretory cells to undergo mucous cell differentiation. Indeed, prior studies have shown that overexpression of active Notch1 intracellular domain in basal stem cells or secretory cells can directly induce mucous cell differentiation^39,42^. Conversely, blocking Notch, including Notch2 and Notch3, has been shown to reduce mucous metaplasia^43–45^. Furthermore, we previously demonstrated that stem cells utilize Jag2 to send a feed-forward signal to secretory cells to affect their fate decision^1^. However, we now demonstrate that the loss of airway basal stem cell-derived Jag2 does not prevent mucous metaplasia following OVA sensitization and challenge (Figure S4M, S4N). To assess whether the loss of basal stem cell-derived Dll1 would alter epithelial intrinsic mechanisms of goblet cell formation, we inhibited Dll1 expression using lentiviral-delivered shRNAs in primary mouse airway basal stem cells in the absence of immune cells in air-liquid interface (ALI) cultures. ALI cultures were treated with recombinant IL-13 to induce mucous cell formation (Figure S6A)^42,44,46,47^. Mucous metaplasia was not altered by decreased expression of Dll1 (Figure S6B, S6C). In addition, overexpression of Dll1 in basal stem cells did not induce mucous cell differentiation (Figure S6D-S6F), indicating that basal stem cell-derived Dll1 is not sufficient, nor is it required for mucous metaplasia. Finally, we tested whether basal stem cells are necessary *in vivo* for IL-13-induced mucous metaplasia in the trachea. Intranasal administration of recombinant IL-13 resulted in extensive mucous metaplasia in Tam-treated CK5-p63 mouse tracheas (Figure S6G-S6I). Therefore, the absence of basal stem cells does not cause an underlying defect in the intrinsic capacity for secretory cells to differentiate into mucous cells *in vivo*. In aggregate, our combined observations lead us to conclude that allergen-induced mucous metaplasia does not require a feed-forward signal from basal stem cells. Thus, in the absence of evidence for a direct epithelial intrinsic stem cell contribution to goblet cell metaplasia and the lack of an effect of stem cell loss on tuft and NE cell populations (which are known to have immunomodulatory effects), our findings all point to an essential role for airway basal stem cells in maintaining a functional pool of IAMs which in turn are important for the development of allergic airway inflammation and mucus metaplasia.

### Notch signaling components are highly expressed in human airway basal stem cells and airway mononuclear phagocytes

To assess whether human basal stem cells possess the capacity to regulate airway mononuclear phagocytes (aMNP) via Notch signaling, we re-analyzed single-cell transcriptomes from baseline and allergen-challenged airway specimens from allergic human subjects with and without asthma. Specifically, scRNAseq profiles, obtained from 21 lower airway brushes collected from 8 human subjects with systemic sensitization to aeroallergens as previously described^48^, were found to contain 8,510 LYZ^+^ aMNP cells that could be clustered into 14 clusters, including DCs, macrophages, and monocyte-derived cells (MCs) (Figure 6A). The DC clusters corresponded to well-defined DC1, DC2, AXL^+^/Siglec6^+^ DC, and plasmacytoid DC subclusters that had been previously characterized using peripheral blood samples^49^. The monocyte-derived clusters (MC1-4) showed expression of classical monocyte markers, including CD14, but otherwise displayed a combination of DC and macrophage markers consistent with known monocyte plasticity. Tissue macrophage subclusters displayed *MAFB*, high levels of complement family genes (*C1QA, C5AR1*)^50–53^, and low levels of ZBTB46 compared to DC populations (Figure 6B). We next evaluated whether any of the aMNP clusters had a similar transcriptomic profile to murine IAM. Discriminatory genes for murine IAMs generated previously^13^ were mapped onto human orthologs to define an IAM signature. Gene set enrichment analysis was performed to evaluate for an enrichment of the IAM signature across all aMNP clusters. Notably, only the macrophage cluster Mac2 showed positive enrichment for IAM marker genes (Figure 6C and Figure S7A, S7B). Mac2 and IAM showed considerable overlap with genes that serve as canonical macrophage markers, including complement genes (*C1q*) and *MAFB* (Figure 6B and Figure S7B). Like murine IAMs, Mac2s express CD11c (*ITGAX*), high MHC II (*HLA-DQB1*), and other genes associated with antigen presentation (*CIITA, CD74*) (Figure 6B and Figure S7B). Mac2 also express the genes for Langerin (CD207) and MerTK (Figure S7B). Additionally, Mac2 and IAMs express *MMP12* and *AXL,* further suggesting similar functional roles involving tissue remodeling and efferocytosis, both implicated in asthma pathobiology (Figure 6B) ^54,55^.

**Figure 6.**
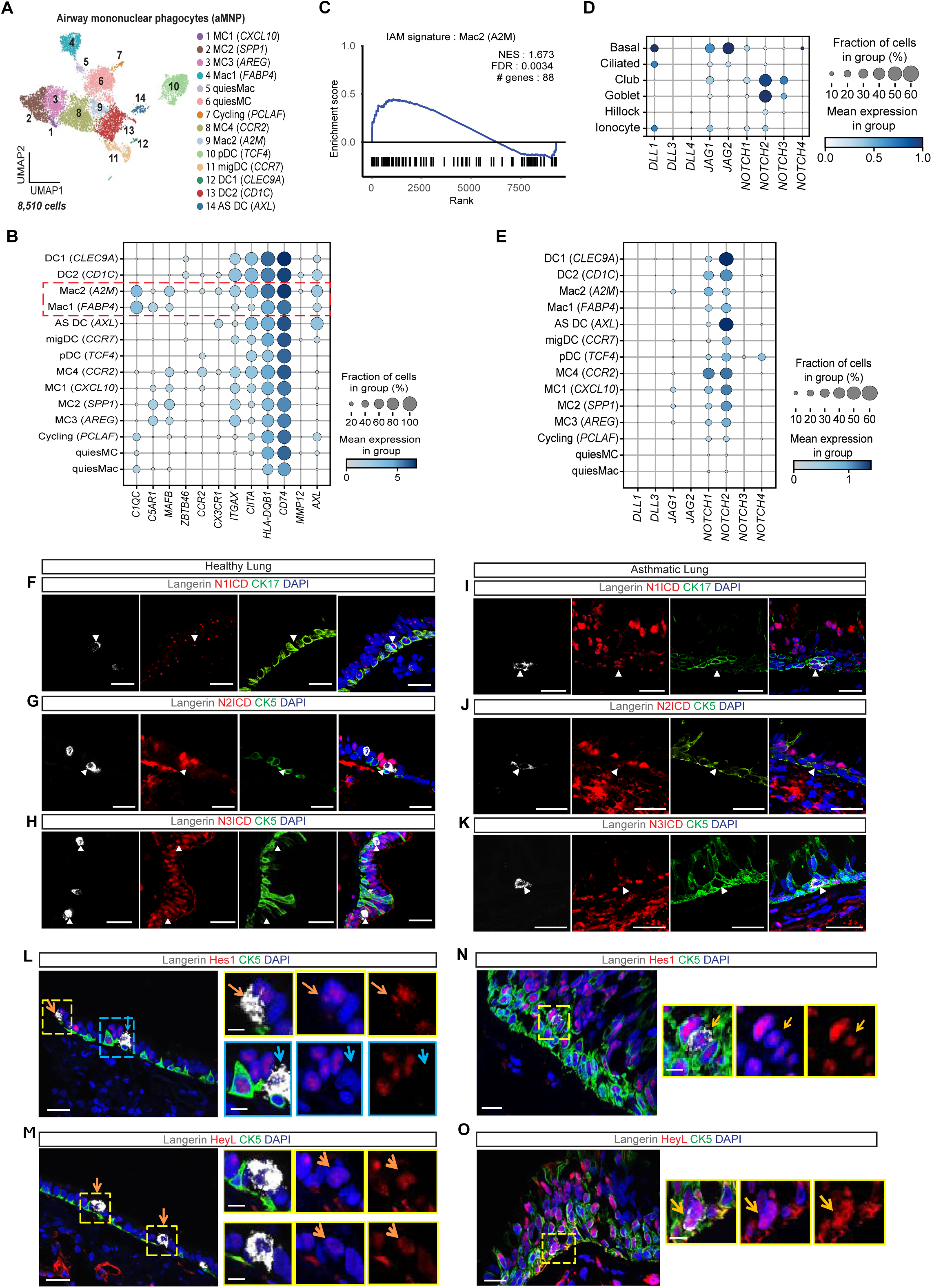
Stem-immune Notch signaling in the human airway epithelium. **A**, UMAP embedding derived from subclustering of 8,510 mononuclear phagocytes (MNP). MC: monocyte-derived cell. Mac: macrophage. DC: dendritic cell. **B,** Dot plot depicting gene expression levels and percentage of cells expressing genes across MNP clusters. The red dashed box highlights macrophage populations Mac1, Mac 2. **C,** Gene set enrichment analysis of Mac2 based on an IAM gene set from Engler *et al* (PMID: 33378665). FDR: false discovery rate. NES: normalized enrichment score. **D, E,** Dot plot depicting Notch ligand and receptor gene expression levels and percentage of cells expressing genes across airway epithelial cell clusters (**D**) and MNP clusters (**E**). **F-K,** Immunostaining for Notch1 intracellular domain (N1ICD) **(F, I),** Notch2 intracellular domain (N2ICD) **(G,J)** or Notch3 intracellular domain (N3ICD) **(H,K)** (red), the intraepithelial immune cell marker Langerin (white) and basal stem cell marker CK5 or CK17 (green) in the human small airway of two healthy donors **(F-H)** or two asthma patients **(J-K).** Arrows indicate Langerin+ cells with Notch activity. **L,M,** Immunostaining for the Notch target genes Hes1 (L) and HeyL (M) (red) with Langerin (white) and CK5 (green) in healthy human airways. Orange arrows point to Hes1+ (L) or HeyL+ (M) Langerin+ cells. Blue arrow points to a Hes1 negative Langerin+ cells (L). **N,O,** Immunostaining for Hes1 (N) and HeyL (O) (red) with Langerin (white) and CK5 (green) in asthmatic human airways. Orange arrows point to Hes1+ (N) or HeyL+ (O) Langerin+ cells. Nuclei stained with DAPI (blue). Scale bars, 20μm. Scale bars in insets in L-O, 5μm. See also Figure S7.

We next analyzed the expression of Notch signaling components within the human aMNP and airway basal stem cell subclusters. Human basal stem cells showed robust expression of Notch ligands *DLL1*, *JAG1*, and *JAG2* (Figure 6D) mirroring the previously described expression patterns in mouse basal cells^1,39^. Correspondingly, we identified strong expression of Notch receptors Notch1 and Notch2 across most aMNP populations (Figure 6E), including Mac2. We could identify Mac2 cells in human airway tissue sections as Langerin^+^ intraepithelial cells that co-express MerTK and C1q (Figure S7C, S7D). Like IAMs, Langerin^+^ Mac2 are found adjacent to and immediately above CK5^+^ and CK17^+^ basal cells (Figure 6F-6H). Subsets of intraepithelial Langerin^+^ Mac2 cells showed evidence of Notch activation *in vivo* based upon nuclear staining for N1ICD, N2ICD, and N3ICD in both healthy and asthmatic human distal airways (Figure 6F-6K, Figure S7E-S7H). Detection of the Notch target genes Hes1 (Figure 6L, 6N) and HeyL (Figure 6M, 6O) in Langerin+ intraepithelial cells confirms Notch activation in Mac2 cells in healthy airways (Figure 6L, 6M) and in asthmatic airways (Figure 6N, 6O). These results suggest that human Mac2 myeloid cells are the human counterparts of murine IAMs since these two cell populations share multiple features including their pattern of gene expression indicative of a specific myeloid subset, their intraepithelial location, their proximity to basal stem cells, and in particular their expression of MHC II-associated genes and evidence of tonic Notch activation.

## DISCUSSION

Here, we show that airway basal stem cells signal to a newly characterized intraepithelial macrophage population (IAMs) and are required for the maintenance of their differentiated state and function (Figure 7). IAMs have a characteristic dendritiform morphology and are localized adjacent to airway basal stem cells. Although prior work has characterized these cells as DCs, our data demonstrate that IAMs are distinct from the Zbtb46 lineage (Figure 1B, S1A, S1F) and possess a unique signature that is aligned with a macrophage population. In addition, IAMs express high levels of MHC II, like other antigen-presenting cells, and are essential for allergic airway inflammation induced by the model antigen OVA in murine proximal airways (Figure S5).

**Figure 7.**
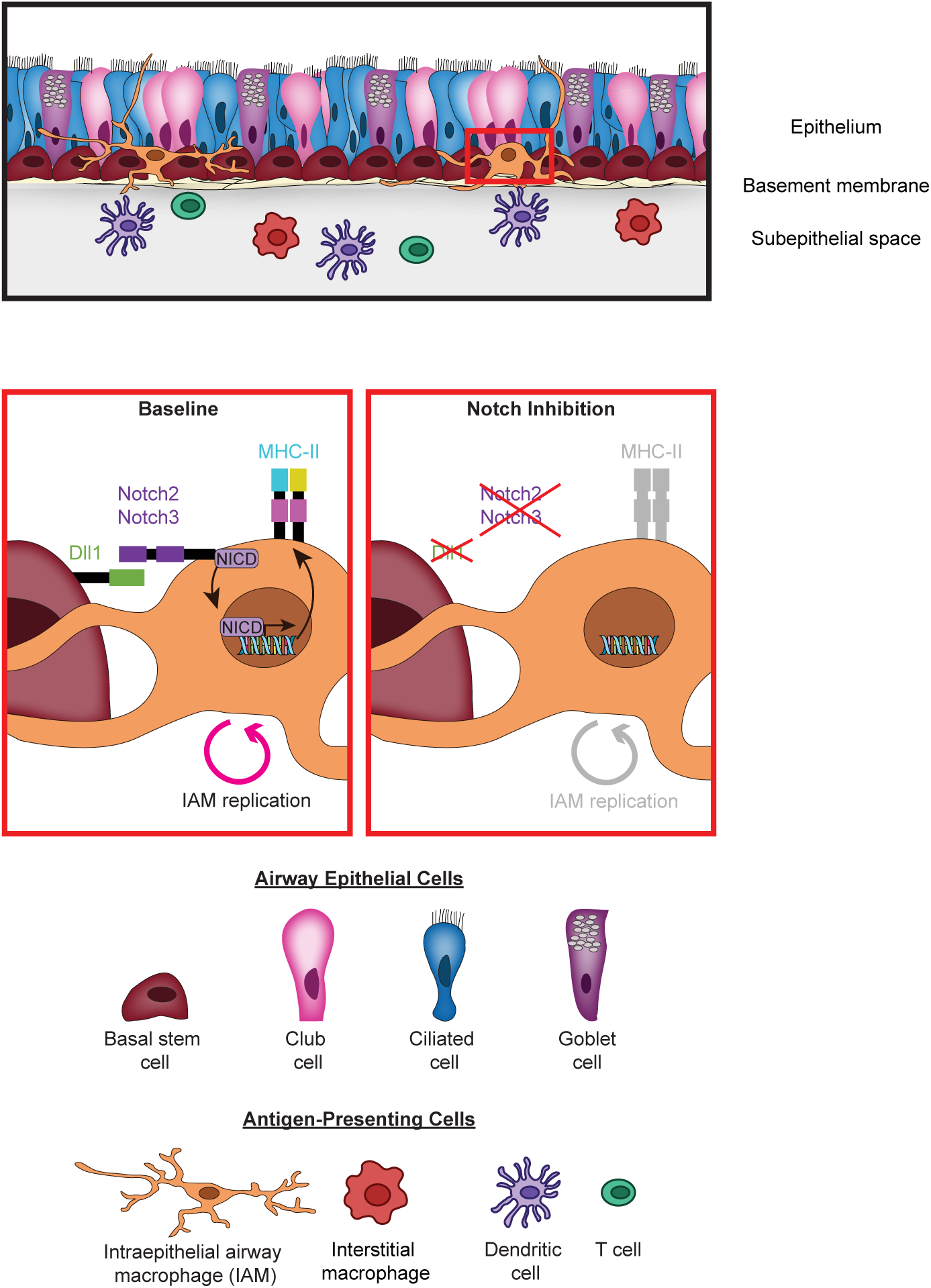
Airway basal stem cells locally regulate the maintenance of intraepithelial airway macrophages (IAMs). Basal stem cell derived-Dll1 activates Notch2 and Notch3 in IAMs to maintain a functional differentiated state. The loss of basal stem cell-derived Dll1, or Notch signaling reception in IAMs results in the loss of MHC II expression and reduces IAM replication.

We demonstrate that airway basal stem cells are required for the maintenance of functional IAMs and that direct cell-to-cell signaling is necessary for MHC II expression. Specifically, airway basal stem cells supply the Notch ligand Dll1, which may engage Notch2 and/or Notch3 receptors on IAMs. Previous work has underscored the interaction of epithelial stem cells and local macrophage populations. Within the alveolus, alveolar type 2 (AT2) epithelial stem cells stimulate and maintain alveolar macrophage maturation through secretion of GM-CSF^56^, and disruption of this interaction results in dysfunctional immature macrophages causing the lung disease pulmonary alveolar proteinosis^57^. In addition, CD200 signaling to alveolar macrophages via CD200R restrains macrophage inflammatory activity ^58^. Thus, in both the proximal airways and alveoli, stem cells are essential for the maintenance of local functional immune cell populations. Interestingly, the functional differentiation of IAMs is largely dependent on direct cell-cell communication with airway basal stem cells, and in contrast to alveolar macrophages have a higher rate of cell turnover (tissue-resident half-life of approximately 6 weeks, Figure 3D) and ongoing recruitment from blood monocytes during homeostasis. Interestingly, mammary gland stem cells have also been shown to maintain macrophages using Notch signaling, potentially pointing to a conserved arrangement of directly communicating cells that allows for coupled regeneration and inflammation in different tissues^37^.

It has been previously shown that MHC II expressing airway macrophages present antigens to airway resident T cells and can transfer antigens to subepithelial DCs that then migrate to draining lymph nodes^25^. This suggests a model wherein IAMs perform local antigen presentation in the airway, guided by direct signals from their surrounding tissue, while DCs propagate responses in draining lymphoid tissues after receiving signals from secreted epithelial cytokines and chemokines.

Loss of Notch signaling in IAMs resulted in decreased airway inflammation following antigen challenge. The reduced allergic inflammation is likely the cause of the reduced goblet cell metaplasia in this model, presumably due to decreased local concentrations of IL-4 and IL-13. We suspect this is a result of impaired antigen presentation to T cells and/or transfer of antigen to airway DCs by IAMs, although this will require further confirmation. The OVA model of asthma has the advantage of minimal direct innate immune cell stimulation as opposed to other models of asthma, such as the house dust mite or Alternaria models, relying largely on antigen-presentation to primed T cells.

Consistent with our studies, prior work has shown that inhibition of Notch receptors with blocking peptides or antibodies impairs allergic airway inflammation in a murine model of asthma^43–45^. In addition, previous studies have highlighted roles for Notch signaling in type 2 inflammation; however, these studies focused on the direct effects of Notch signaling on T cells, affecting their polarization and egress from lymph nodes^59–62^. In contrast, our findings demonstrate a novel role for tonic Notch signaling between airway basal stem cells and IAM, suggesting a role for basal stem cells as an immune cell niche. Interestingly, the effects on allergic airway inflammation were isolated to the portion of the murine airways with pseudostratified epithelium containing basal stem cells (trachea and main bronchi), suggesting compartmentalization of the inflammatory response and different mechanisms of airway inflammation depending on the structure of the airway epithelium. Given that asthma pathology in humans is mostly localized to airways with pseudostratified epithelium, we believe that these findings are highly relevant for asthma pathogenesis. We speculate that the direct functional coupling of local stem cell-mediated regeneration and immune responses helps to promote a compartmentalized inflammatory response to allergic stimuli.

In addition, for the first time, we identify a class of human intraepithelial airway HLA-positive macrophages (Mac2) that resemble murine IAMs, being located adjacent to basal stem cells and exhibiting Notch activation. This suggests a similar direct stem-immune interaction is required for the maintenance and function of these macrophages in human airways. Altogether, our data suggest that manipulation of airway basal stem cell-IAM cross-talk could be used therapeutically to regulate immune responses in the airway.

### Limitations of the study

We identify and further characterize a recently described novel immune cell population that is defined by its location (intraepithelial), morphology (dendritiform), and unique transcriptional signature. Since myeloid cells express many similar markers, a major limitation has been to find a unique Cre driver line that would target only IAMs without impacting other macrophage or DC populations. In the future, further characterization of these cells to identify unique markers will help develop a better strategy to specifically target IAMs.

In addition, based on the nature of the niche signal that requires physical cell-cell contact, we have pointed out the importance of local stem-immune interactions, clearly demonstrating the compartmentalization of the response. This limits the use of whole tissue analysis and necessitates image-based analysis.

The study of tonic Notch activation offers several challenges due to the low levels of expression of key Notch compenents. In fact, signal amplification is necessary to detect NICDs by antibody staining. Additionally, scRNASeq does not capture the expression of specific Notch components, including Notch target genes, and Notch3, whose expression is generally lower than Notch1 and Notch2^1^. More sensitive methods should be implemented to capture low levels of gene expression, and protocols must be optimized to detect low-level signals.

Our work shows evidence of an IAM-like cell population in human airways. Further investigation is required to demonstrate stem-derived niche signals to IAM-like cells in the human airway epithelium. Although a major obstacle is the low number of IAMs limiting their isolation and culture, optimization of *in vitro* models to co-culture both cell types will be helpful in unraveling their intercellular interactions and evaluating the relevance of targeting stem-immune direct communication as a therapeutic option in disease.

## Acknowledgments

We thank Adam Glick for providing the *CK5rtTA* mice, Brigid Hogan for providing *CK5-creER* mice, Alea A. Mills for kindly sharing the *p63 floxed* mice, Rafi Kopan for the *Dll1 floxed* mice and Yibin Kang and Hans Clevers for the *Dll1-mCherry* mice. We also thank Barry Stripp for providing the goat anti-SCGB1A1 antibody. We wish to extend our thanks to all of the current and former members of the Rajagopal Laboratory, the Medoff Laboratory, the Pardo-Saganta Lab, people involved in the maintenance of mouse colonies, the HSCI flow cytometry core facility and the flow core at CIMA, as well as the histology core at MGH, and all the colleagues that spent a minute providing their feedback, especially Carrie Sokol and Mikael Pittet for their comments, guidance and help in the presentation of the results.

This research was supported by the New York Stem Cell Foundation (J.R. is a New York Stem Cell Foundation-Robertson Investigator), by a National Institutes of Health-National Heart, Lung, and Blood Institute Early Career Research New Faculty (P30) award (5P30HL101287-02), RO1s (RO1HL118185 and RO1HL164563) from NIH-NHLBI (to J.R.), a Harvard Stem Cell Institute (HSCI) Junior Investigator Grant (to J.R.), a grant from the Department of Defense (W81XWH) and a Sanofi iAward to B.D.M. an NIH-NHLBI grant T32HL116275 and American Thoracic Society Award to T.K., a grant from Instituto de Salud Carlos III ISCIII “PI17/01346”, co-funded by ERDF,“A way to make Europe” (to B.S.), an AEFAT award (to B.S.), an AECC Junior Investigator Grant exp. AIO16163636SAEZ (to B.S.), a Ramon y Cajal Award RYC-2015-18580 co-founded by FSE (to A.P-S.), a grant from Gobierno de España SAF2017-89908-R (MINECO/AEI/FEDER, UE.) (to A.P-S), and a grant from Gobierno de Navarra GNS80/2016 co-founded by FEDER (to A.P-S).

## Author contribution

T.G.K. designed and performed the experiments, analyzed and interpreted the data, optimized protocols for wholemount staining and imaging of tracheal explants and edited the manuscript; B.S. and M.R. performed *in vivo* experiments and analysis, contributed to the *in vitro* experiments, helped with the interpretation of the data and edited the manuscript; A.E-Z and D.L. processed samples, performed immunostaining analysis and generated final images; D.A., N.N., J.V. and J.G. helped with *in vivo* experiments; E.P., L.V. and Z. B-I processed samples from *in vivo* experiments, optimized immunostaining protocols, generated tools and helped with *in vitro* experiments; N.P.S., A-C.V., J.A., Y.Z. and A.H., re-analyzed scRNASeq datasets; V.V. optimized the detection of intraepithelial myeloid cells on tracheal explants; M.S. analyzed NE cells; S.L.S. generated cartoons, reorganized figures and helped editing the manuscript; M.G-C. contributed to the analysis of the phenotype of CK5-p63 mice; H.M. and A.W. provided protocols and reagents and helped perform the *in vitro* experiments; B.L., A.P. and K.Y. organized, reestablished and provided different mouse lines; P.R.T. and R.Z. contributed to the *in vivo* experiments using the mouse model of basal cell depletion; B.C. helped with the analysis of the inflammatory response; J.Z, F.P. and J.L.C. contributed to the interpretation of the data and provided input on the manuscript; J.R. and B.D.M. co-designed the study, supervised the project and co-wrote the manuscript; A.P-S. conceived the study, designed and performed the experiments, analyzed and interpreted the data, supervised the project, coordinated the entire team, and wrote the manuscript.

## Declaration of interests

The authors declare no competing interests.

**Correspondence and requests for materials** should be addressed to Ana Pardo-Saganta or Benjamin D. Medoff.

**Supplementary Information** is available for this paper, it includes Supplementary Figures and is provided as a separate document.

## Data Availability

The following publicly available datasets in Gene Expression Omnibus under accession numbers GSE103354 ^2^ and GSE193816 ^48^ were reanalyzed.

## Code availability

Codes for analysis of scRNASeq data are available at GitHub (https://github.com/tristankooistra/NotchAirwayMacs)

## Methods Summary

*CK5-rtTA*^1^, *tet(O)cre* (JAX 006224), *CK5-creER*^1^, *LSL-YFP* (JAX 006148), *TdTomato* (JAX7909), *Zbtb46-Cre* (JAX028538), mTmG (JAX007576), *CD11c-Cre* (JAX008068), *p63 ^fl/fl^* ^29^, *Jag2 ^fl/fl^* ^1^, *Dll1^fl/fl^* ^63^, *Dll1-mCherry* ^38^, CD11c-mCherry ^64^, *Rosa DTA* (JAX 009669), *Cx3cr1-GFP;Ccr2-RFP* (JAX 032127), *Cx3cr1-CreER* (JAX 021160), *RBPJk ^fl/fl^*^1^, *CBF:H2B-Venus* (JAX 020942) and C57BL6/J (JAX 000664) mice were previously described. All procedures and protocols were approved by the Ethics and Animal Care Committee. Human lung tissue was obtained from explanted lungs through the DZL Biobank Giessen in compliance with current regulation. Informed consent was obtained from each subject and all protocols were approved by the Ethical Committee. Tissue preparation, immunofluorescence, microscopy, cell sorting, quantitative RT-PCR, ELISA analysis and basal cell culture were performed using previously described protocols^1,29,34,41,46^. Cell counts were performed on tracheal sections and on wholemounts analyzing multiple fields per sample (mouse trachea). scRNA-seq analysis were performed using previously published datasets^2,48^.

## Statistical analysis

The standard error of the mean was calculated from the average of the indicated number of samples in each case (n= biological replicates / condition / experiment). All the experiments were repeated two or three times. The number of samples per condition is indicated in each case in the figure legend. Multiple 20x or 40x magnification fields per sample were counted and is indicated in detail in Extended Methods. No animals were excluded from the analysis. Data was analyzed and compared among groups by Student’s t-test (unpaired, two-tailed test) or two-way ANOVA. In the experiments where multiple fields per trachea were counted, the data was analyzed by nested t test. A *p*-value of less than 0.05 was considered significant. The analysis was performed with GraphPad Prism 10.0 software.

